# Effective population size and durability of plant resistances in the potato cyst nematode *Globodera pallida*

**DOI:** 10.1101/223826

**Authors:** Josselin Montarry, Eric J. Petit, Sylvie Bardou-Valette, Romain Mabon, Pierre-Loup Jan, Sylvain Fournet, Eric Grenier

## Abstract

- The effective size of a population is the size of an ideal population which would drift at the same rate as the real population. The balance between selection and genetic drift depends on the population size expressed as the genetically effective population size (N_e_), rather than the real numbers of individuals in the population (N).
- The objectives of the present study were to estimate N_e_ in the potato cyst nematode *Globodera pallida* using artificial populations and to explore the link between N_e_ and the durability of plant resistances.
- Using a temporal method on 24 independent pairs of initial and final populations, the median N_e_ was 58 individuals.
- N_e_ is commonly lower than N but in our case the N_e_/N ratio was extremely low because *G. pallida* populations deviate in structure from the assumptions of the ideal population by having unequal sex-ratios, high levels of inbreeding and a high variance in family sizes. The consequences of a low N_e_ could be important for the control of phytoparasitic nematodes because *G. pallida* populations will have a low capacity to adapt to changing environments unless selection intensity is very strong, which could be greatly beneficial for long-term use of plant resistances.

## Introduction

Mutation, migration, selection and genetic drift determine the evolution of populations, but genetic drift has a much greater impact and selection is less effective in small than in large populations (Frankham *et al*., 2002). When both factors are operating, selection (deterministic) predominates in large populations, while genetic drift (stochastic) predominates in small populations (Kimura *et al*., 1963; Nei *et al*., 1975; Gherman *et al*., 2007). Indeed, within small populations, the random sampling of gametes due to genetic drift leads to i) random changes in allele frequencies from one generation to the next, ii) loss of genetic diversity and fixation of alleles within populations, and consequently to iii) rapid diversification among fragmented populations from the same original source. The balance between selection and genetic drift depends on the population size expressed as the genetically effective population size (N_e_), rather than the real numbers of individuals in the population (N, the census size). The effective size of a population is the size of an ideal population which would drift at the same rate as the observed population (Wright, 1931). Thus, the N_e_ of a population is a measure of its genetic behaviour, relative to that of an ideal population, characterized by no migration, no mutation, no selection, no overlapping generations, equal sex-ratios, constant size in successive generations (i.e. on average one offspring per adult), random union of gametes, and a Poisson distribution of family sizes (Frankham *et al*., 2002). Any characteristic of the real population that deviates from those characteristics of the ideal population will cause the effective size (Ne) to differ from the number of individuals in the population (N).

Plant pathogens impose a major constraint on food production worldwide. They are often combated with pesticides, but the need to develop more sustainable production systems fuels a trend towards a limitation of pesticide applications. Among possible alternatives to chemical control, plant resistances look promising, but can only be used in the long term and accepted in the short term if their durability is ascertained. Durability of host resistance was defined as the persistence of resistance efficiency when resistant cultivars are used over long periods, on large surfaces and in the presence of the target pathogen (Johnson, 1981, 1984): it therefore depends primarily on the rhythm of adaptive changes affecting pathogen populations in response to selection by host resistance. The speed of fixation of an advantageous allele depends on the difference in its selection coefficient from that of the other alleles (ΔS) but also on the action of genetic drift, which is influenced by the effective population size (N_e_). The value of the product N_e_ * ΔS determines whether the population will be mainly under the influence of selection or of genetic drift (Crow & Kimura, 1970; Fraser, 1972; Charlesworth, 2009). Consequently, the selection of virulent alleles (the virulence being defined as the ability to infect a resistant host – Vanderplank, 1963; Gandon & Michalakis, 2002; Tellier & Brown, 2007, 2009) by resistant plants could be partly compromised by a low effective population size.

The effective size has been widely investigated both theoretically (Crow & Kimura, 1972; Nei & Tajima, 1981; Tajima & Nei, 1984; Criscione & Blouin, 2005) and experimentally (Johnson *et al*., 2004; Wang, 2005; Araki *et al*., 2007) in a broad variety of organisms. For plant pathogens, the effective size is a very important point to take into account in order to manage plant resistances (e.g. Fabre *et al*., 2012). N_e_, and thus the importance of genetic drift, has been explored for several plant viruses (e.g. Betancourt *et al*., 2008; Monsion *et al*., 2008; Zwart *et al*., 2011; Gutiérrez *et al*., 2012; Fabre *et al*., 2012, 2014) and fungi (e.g. Damgaard & Giese, 1996; Zhan *et al*., 2001; Duan *et al*., 2010; Stukenbrock *et al*., 2011). Regarding plant parasitic nematodes there is only one recent study exploring the N_e_ of the beet cyst nematode *Heterodera schachtii* from wild populations, sampled on *Beta maritima* (Jan *et al*., 2016). The objectives of the present study were to estimate the N_e_ of the potato cyst nematode *Globodera pallida*, and to explore the link between N_e_ and the durability of plant resistances.

Several methods are dedicated to estimate effective population sizes (Wang, 2005; Palstra & Ruzzante, 2008; Luikart *et al*., 2010). Methods using a single-sample estimate N_e_ from the linkage disequilibrium and/or the heterozygote excess (Pudovkin *et al*., 1996; Tallmon *et al*., 2008; Waples & Do, 2008), whereas temporal methods estimate N_e_ from the variation of allelic frequencies between two temporally spaced samples. Deviations from Hardy-Weinberg equilibrium due to heterozygote deficits have been recorded for three plant parasitic nematode species (*Globodera pallida* - Picard *et al*., 2004, *Heterodera schachtii* - Plantard & Porte, 2004, and *Globodera tabacum* - Alenda *et al*., 2014) and recently attributed to both consanguinity and sub-structure at the plant scale (Montarry *et al*., 2015). Those biological characteristics, inbreeding and sub-structure (Wahlund effect), are known to bias single-sample estimators of N_e_ (Zdhanova & Pudovkin, 2008; Waples & Do, 2010; Holleley *et al*., 2014). Therefore, we estimated N_e_ of *G. pallida* populations by using a temporal method developed by Wang (2001).

Rather than working with natural populations, which could sometimes harbor very low genetic diversity leading to infinite estimated Ne, we decided here to work with artificial populations in order to maximize the allelic diversity in the initial populations and thus to better follow the variation of allelic frequencies between initial and final *G. pallida* populations.

## Materials and Methods

### Biology of *Globodera pallida*

Nematodes are a group of worms that include free-living species such as *Caenorhabditis elegans* as well as many parasitic species of animals and plants. Plant-parasitic nematodes are major parasites that cause considerable economic losses in agriculture: worldwide crop losses caused by nematodes have been estimated around $100 billion per year (Sasser & Freckman, 1987). *Globodera pallida* is a gonochoristic diploid organism with obligatory sexual reproduction, which achieves one generation per year (Adams *et al*., 1982). *G. pallida* is probably native to the Andean Cordillera (Grenier *et al*., 2010), the origin of its unique host genus *Solanum* (Hijmans & Spooner, 2001). This obligate, sedentary endoparasite enters the plant roots as second-stage juveniles (J2) and establishes a specialized feeding structure, the syncytium (Jones & Northcote, 1972), which is a severe nutrient sink for the plant. Sex is environmentally determined through the size of the syncytium that is induced (Sobczak & Golinowski, 2011). Adult males leave the root in order to mate females. The females continue to feed and when egg development is completed, they die and form a cyst, enclosing hundreds of eggs, which constitute a survival stage that can stay viable for several years in soils.

### Initial nematode populations

Twenty-four initial *G. pallida* populations, each composed of 100 cysts, were artificially made by mixing four Peruvian populations genetically differentiated and showing high allelic richness (Otuzco, Peru_252, Peru_286 and Peru_298; Picard *et al*., 2004). Before mixing those populations, the number of larvae was scored for 12 randomly chosen cysts, and a one-way ANOVA showed no significant difference for the number of larvae per cyst between those four populations (*F*_3,44_ = 1.59; *P* = 0.21; Fig. 1). The initial census size was thus estimated by multiplying the number of cysts (i.e. 100) by the mean number of larvae per cyst (i.e. 132, Fig. 1).

**Fig. 1.**
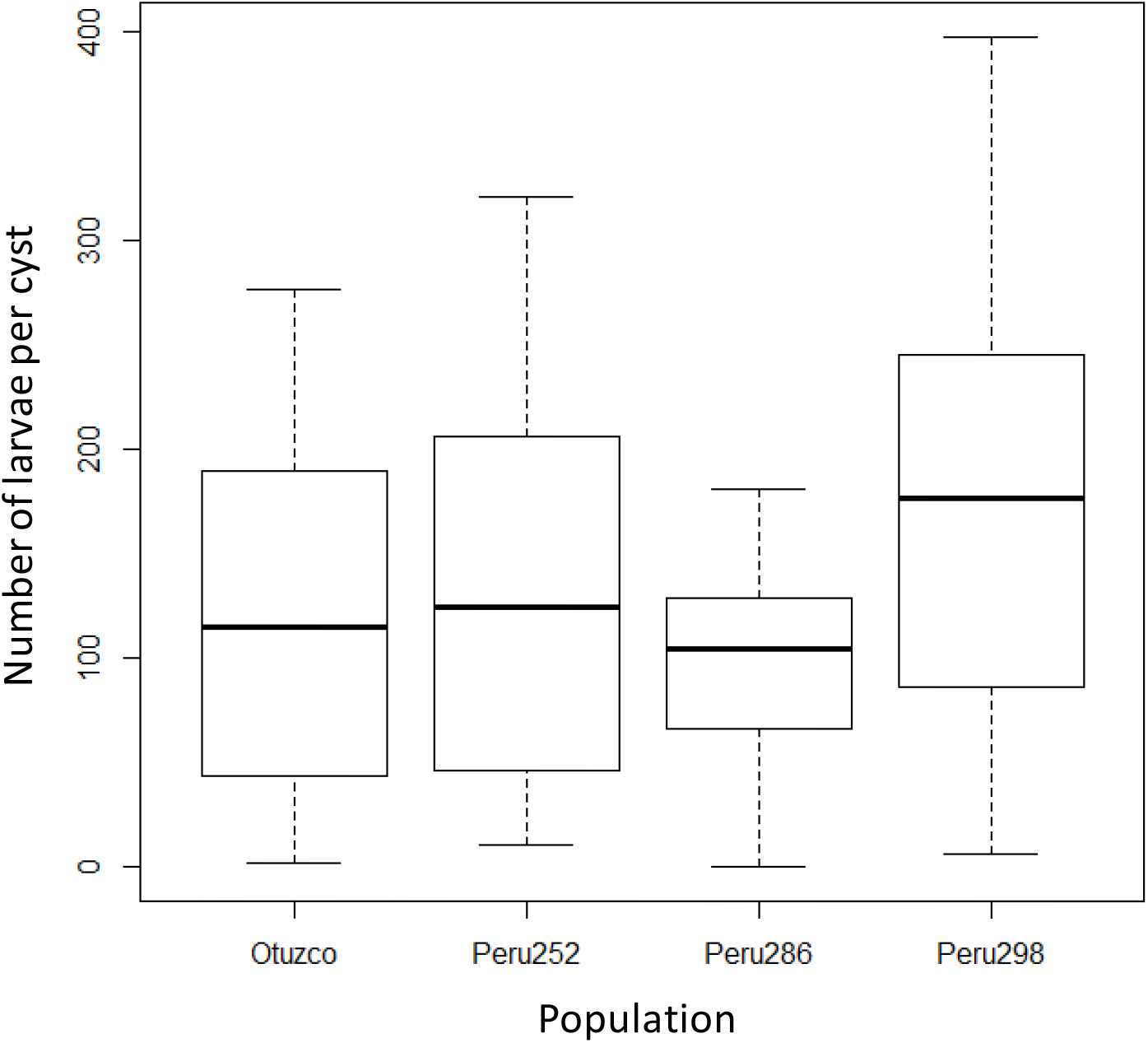
Number of larvae per cyst for the four *G. pallida* Peruvian populations (Otuzco, Peru_252, Peru_286 and Peru_298). No significant difference was observed between those populations (F_3,44_ = 1.59; *P* = 0.21).

We mixed different numbers of cysts from the different Peruvian populations to start with allelic frequencies that differ between initial populations. Each of seven different cyst proportions was replicated three times, apart from the equal mix, which was replicated six times, for a total of 24 initial populations (Table 1). Seven initial populations (Pi_A to Pi_G), composed of 50 cysts, were also prepared in the same proportions for the estimation of initial allelic frequencies.

**Table 1.**
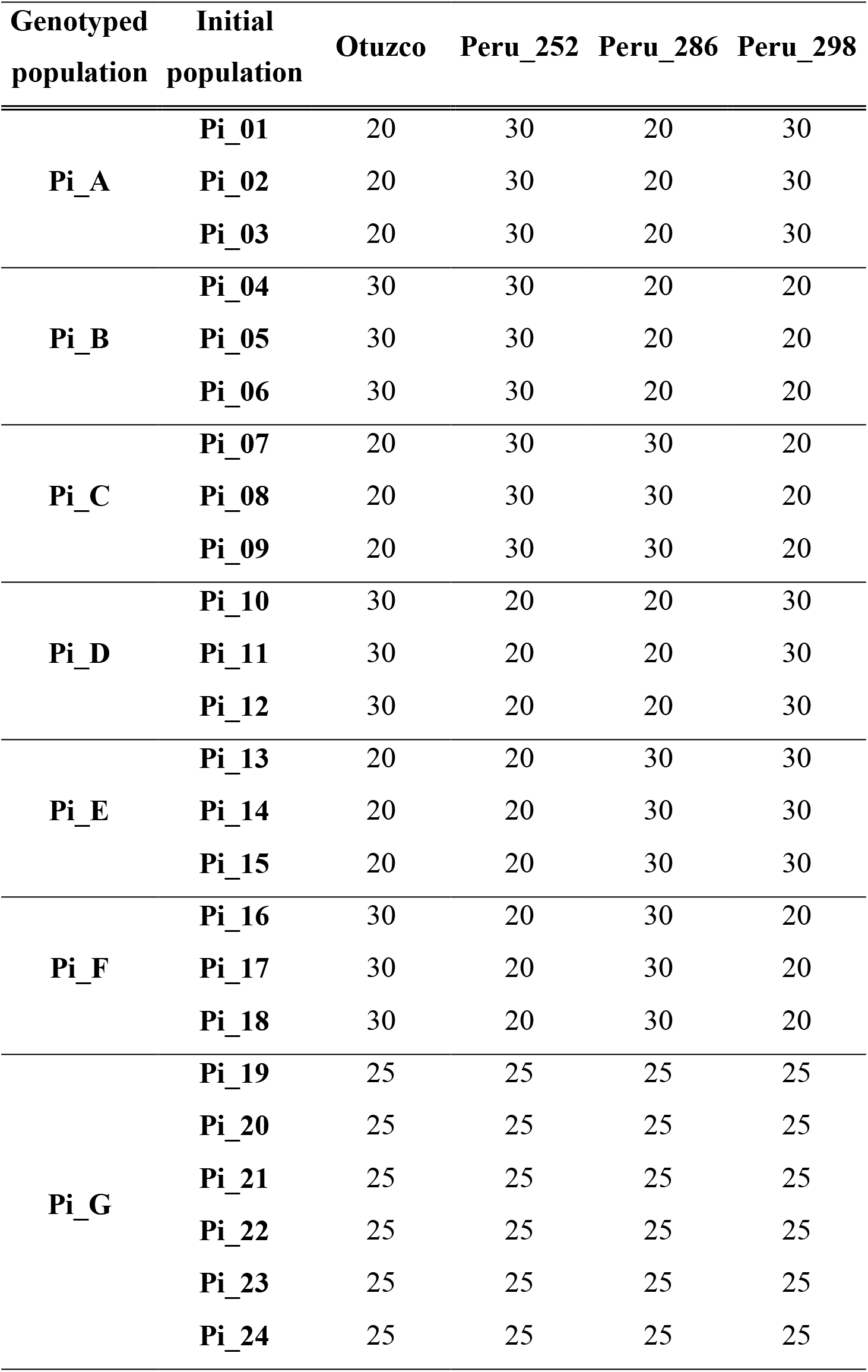
Number of cysts from each of the four Peruvian *G. pallida* populations (Otuzco, Peru_252, Peru_286 and Peru_298) used to construct the 24 initial populations (Pi_01 to Pi_24). The estimation of the initial allelic frequencies was performed through the genotyping of the seven initial populations (Pi_A to Pi_G) composed of 50 cysts mixed using the same proportions.

| Genotyped population | Initial population | Otuzco | Peru_252 | Peru_286 | Peru_298 |
| --- | --- | --- | --- | --- | --- |
| Pi_A | Pi_01 | 20 | 30 | 20 | 30 |
|  | Pi_02 | 20 | 30 | 20 | 30 |
|  | Pi_03 | 20 | 30 | 20 | 30 |
| Pi_B | Pi_04 | 30 | 30 | 20 | 20 |
|  | Pi_05 | 30 | 30 | 20 | 20 |
|  | Pi_06 | 30 | 30 | 20 | 20 |
| Pi_C | Pi_07 | 20 | 30 | 30 | 20 |
|  | Pi_08 | 20 | 30 | 30 | 20 |
|  | Pi_09 | 20 | 30 | 30 | 20 |
| Pi_D | Pi_10 | 30 | 20 | 20 | 30 |
|  | Pi_11 | 30 | 20 | 20 | 30 |
|  | Pi_12 | 30 | 20 | 20 | 30 |
| Pi_E | Pi_13 | 20 | 20 | 30 | 30 |
|  | Pi_14 | 20 | 20 | 30 | 30 |
|  | Pi_15 | 20 | 20 | 30 | 30 |
| Pi_F | Pi_16 | 30 | 20 | 30 | 20 |
|  | Pi_17 | 30 | 20 | 30 | 20 |
|  | Pi_18 | 30 | 20 | 30 | 20 |
| Pi_G | Pi_19 | 25 | 25 | 25 | 25 |
|  | Pi_20 | 25 | 25 | 25 | 25 |
|  | Pi_21 | 25 | 25 | 25 | 25 |
|  | Pi_22 | 25 | 25 | 25 | 25 |
|  | Pi_23 | 25 | 25 | 25 | 25 |
|  | Pi_24 | 25 | 25 | 25 | 25 |

### Final nematode populations

The 24 initial *G. pallida* populations were inoculated to 24 potato plants of the susceptible potato cultivar Désirée. Because Désirée is a susceptible cultivar, there is no *a priori* reason to expect directional selection in favour of an allele. Moreover, there is also no *a priori* reason to assume that any selection affecting the allelic frequencies is acting across plants (e.g. favouring an allele in a plant and selecting against it in another plant). We have thus assumed that selection is negligible.

For each initial population, the 100 cysts were locked in a tulle bag and placed in a 13cm pot filled with a soil mixture free of cysts (2/3 sand and 1/3 natural field soil). Tubers were then planted and covered with the same soil mixture. Plants were grown during four months in a climatic chamber regulated at 20°C with a 16h photoperiod. During that period, the monovoltine species *G. pallida* achieved only one generation. Newly formed cysts from the 24 final populations were then extracted from the soil using a Kort elutriator and stored at 4°C before genotyping. The number of newly formed cysts was counted for each final population and the number of larvae per cyst was scored for 12 randomly chosen cysts for seven final populations among the 24 (i.e. one randomly chosen population per initial proportion). The final census size was thus estimated by multiplying the mean number of cysts by the mean number of larvae per cyst.

### Microsatellite genotyping

The 31 *G. pallida* populations (i.e. seven initial and 24 final populations) were genotyped using 12 neutral microsatellite markers (Gp106, Gp108, Gp109, Gp111, Gp112, Gp116, Gp117, Gp118, Gp122, Gp126, Gp135 and Gp145) developed by Montarry *et al*. (2015) directly from the *G. pallida* genome (Cotton *et al*., 2014). For each population, from 26 (for Pi_E) to 40 (for Pi_C) larvae, coming from distinct and randomly chosen cysts, were successfully genotyped. Two multiplex panels were used to reduce the time and cost required to genotype the 1,105 individuals at the 12 loci.

DNA from single larva (i.e. one second-stage juvenile J2) was extracted following a procedure using sodium hydroxide and proteinase K (Boucher *et al*., 2013). PCR was performed using a 384-well reaction module (BIO-RAD C1000) in a 5 μL volume containing 1X of Type-it Microsatellite PCR kit, 0.4 μM of primer mix and 1 μL of template DNA. Cycling conditions included an initial denaturation at 95 °C for 5 min, followed by 30 cycles of denaturation at 95 °C for 30 s, annealing at 57 °C for 90 s and extension at 72 °C for 30 s, followed by a final extension at 60 °C for 30 min. PCR products were then diluted to 1:25 in sterile water and 3 μL of this dilution were mixed with 0.05 μL of GeneScan 500 LIZ Size Standard (Applied Biosystems) and 5 μL of formamide (Applied Biosystems). Analyses of PCR products were conducted on ABI Prism^®^ 3130xl sequencer (Applied Biosystems). Allele sizes were determined by the automatic calling and binning module of GeneMapper v4.1 (Applied Biosystems) with manual examination of irregular results. To minimize the rate of genotyping errors, a second round of PCR and electrophoresis was performed for 10% of the global number of individuals.

### Population genetic characteristics

Genetic diversity of each nematode population was estimated through allelic richness (A_r_) and unbiased gene diversity (H_nb_) (Nei, 1978). Departure from Hardy-Weinberg equilibrium was tested through the *F*_IS_ estimation for each population. H_nb_ and *F*_IS_ were computed using genetix 4.05.2 (Belkhir *et al*., 1996–2004). The statistical significance of *F*_IS_ values for each population was tested using the allelic permutation method (1,000 permutations) implemented in genetix. A_r_ was estimated on a reduced sample of 26 individuals using the rarefaction method implemented in Populations 1.2.32 (Langella, 2000).

Because heterozygote deficits in cyst nematodes could be due to a Wahlund effect (i.e. sub-structure) and/or to consanguinity (Montarry *et al*., 2015), we used the method of Overall and Nichols (2001) in order to calculate a likelihood surface for the genetic correlation due to population subdivision (*θ*) and the proportion of the population practicing consanguinity (C). The method, which is based on the argument that consanguinity and sub-structure generate distinctive patterns of homozygosity in multilocus data, was applied with a degree of relatedness of 1/4 (see Montarry *et al*., 2015) to all initial and final populations showing significant heterozygote deficits. The most likely parameter combination was identified over a grid of 10,000 combinations of *θ* and C values, and graphs of the likelihood surface were drawn for each nematode population using the statistical software R version 3.1.1 (R Core Team, 2014).

### Effective population size estimation

The temporal method developed by Wang (2001) was used to estimate N_e_ from the 24 independent pairs of initial and final populations of *G. pallida*. That likelihood-based method has been implemented in the MLNE 1.0 software (Wang & Whitlock, 2003).

The effect of initial populations, differing by the proportion of cysts coming from the different Peruvian populations, on N_e_ was tested using an ANOVA. Normality and homogeneity of variances were checked with the Shapiro-Wilk and the Levene tests, respectively, and mean values were compared with a Tukey test (α = 0.05). The correlation between the N_e_ estimates and the number of newly formed cysts in each final population was tested using the Pearson’s correlation test. All statistical analyses were performed using R.

### Exploring the link between N_e_ and resistance durability

The effective population size could be important for the control of plant pathogens because the effectiveness of selection is directly linked to the effective population size: selection is less effective in small than in large populations. Indeed, the probability of fixation of an allele in a population depends on its initial frequency, its selective advantage (or disadvantage) and the effective population size. Kimura (1962) showed that the probability of fixation of allele A is given by:

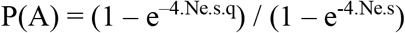

where q is the initial frequency of allele A, s the selection coefficient and N_e_ the effective population size. For a neutral allele, the fixation probability equals its frequency in the population (because genetic drift favours neither allele) but for advantageous alleles, the relationship between the fixation probability and the selection coefficient depends on the effective population size.

Fixing the initial frequency of allele A at 1% (q = 0.01), the relationship between the fixation probability and the selection coefficient was explored for five N_e_ values, i.e. for the estimated N_e_ and for N_e_ two-, five-, 10- or 50-times higher.

Moreover, the selection coefficients exerted by four resistant potato genotypes were estimated using data from Fournet *et al*. (2013), who used these genotypes (96F.376.16, 94T.146.52, 360.96.21 and 60.96.1) to perform an experimental evolution during eight *G. pallida* generations. The selection coefficient *S* was calculated using the univariate breeder’s equation (Lush, 1937): *R* = *h*^2^ * *S*, where *R* is the change in mean phenotype between two generations, *h*^2^ is the heritability, i.e. the proportion of phenotypic variance in the trait that is attributable to genetic effects. Making the assumption that the heritability of the measured trait (the number of produced females), which strongly covaries with fitness, is high, i.e. 0.8 < *h*^2^ < 1, resulted in a range of values for *S* for each potato genotype.

## Results

### Genetic characteristics of initial and final populations

As expected, genetic diversity was high for all initial populations (0.64 < H_nb_ < 0.69 and 6.25 < A_r_ < 7.34; Table 2) and that diversity was conserved from one generation to the next, i.e. for the final populations (0.59 < H_nb_ < 0.70 and 5.54 < A_r_ < 7.14; Table 2). All populations, except one (Pf_05), showed a significant heterozygote deficit, with *F*_IS_ ranging from 0.32 to 0.43 for initial populations and from 0.05 to 0.22 for final populations (Table 2). Those heterozygote deficits were due to consanguinity and substructure for the seven initial populations (Table 2 and Fig S1-A, Supporting Information) and only to consanguinity for the 23 final populations showing significant heterozygote deficits (Table 2 and Fig S1-B, Supporting Information).

**Table 2.** Number of cysts (cysts), number of genotyped individuals (n), genetic diversity (Hnb and Ar) and departure from Hardy-Weinberg equilibrium (*F*_IS_) for each *G. pallida* population (i.e. seven artificial initial populations and 24 final populations). *F*_IS_ values significantly different to zero are indicated in bold. For each population showing a significant heterozygote deficit, *θ* and C values corresponding to the maximum-likelihood were indicated.

| Population | cysts | n | $H_{nb}$ | $A_r$ | $F_{IS}$ | $\theta$ | C |
| --- | --- | --- | --- | --- | --- | --- | --- |
| **Initial populations** |  |  |  |  |  |  |  |
| Pi_A | 100 | 39 | 0.68 | 7.23 | **0.37** | 0.31 | 0.13 |
| Pi_B | 100 | 38 | 0.68 | 7.23 | **0.41** | 0.35 | 0.23 |
| Pi_C | 100 | 40 | 0.65 | 7.34 | **0.32** | 0.25 | 0.24 |
| Pi_D | 100 | 39 | 0.64 | 6.25 | **0.43** | 0.38 | 0.28 |
| Pi_E | 100 | 26 | 0.65 | 6.57 | **0.37** | 0.23 | 0.60 |
| Pi_F | 100 | 38 | 0.69 | 7.30 | **0.39** | 0.25 | 0.60 |
| Pi_G | 100 | 36 | 0.68 | 6.90 | **0.43** | 0.26 | 0.85 |
| **Final populations** |  |  |  |  |  |  |  |
| Pf_01 | 2704 | 28 | 0.65 | 7.00 | **0.14** | 0.00 | 0.52 |
| Pf_02 | 3407 | 33 | 0.66 | 6.51 | **0.14** | 0.00 | 0.45 |
| Pf_03 | 3136 | 38 | 0.68 | 6.31 | **0.16** | 0.00 | 0.54 |
| Pf_04 | 2227 | 36 | 0.59 | 6.35 | **0.11** | 0.03 | 0.35 |
| Pf_05 | 2541 | 34 | 0.60 | 5.54 | -0.01 | . | . |
| Pf_06 | 2082 | 36 | 0.62 | 6.24 | **0.13** | 0.00 | 0.48 |
| Pf_07 | 2053 | 37 | 0.67 | 7.14 | **0.10** | 0.02 | 0.38 |
| Pf_08 | 2511 | 38 | 0.70 | 6.87 | **0.17** | 0.00 | 0.56 |
| Pf_09 | 1459 | 32 | 0.65 | 6.59 | **0.12** | 0.00 | 0.46 |
| Pf_10 | 2193 | 34 | 0.66 | 6.78 | **0.18** | 0.02 | 0.59 |
| Pf_11 | 2425 | 37 | 0.63 | 6.27 | **0.05** | 0.00 | 0.25 |
| Pf_12 | 1749 | 39 | 0.64 | 6.40 | **0.22** | 0.05 | 0.57 |
| Pf_13 | 2893 | 33 | 0.63 | 6.29 | **0.05** | 0.00 | 0.18 |
| Pf_14 | 1391 | 36 | 0.61 | 6.45 | **0.13** | 0.00 | 0.49 |
| Pf_15 | 1060 | 35 | 0.68 | 6.77 | **0.07** | 0.00 | 0.26 |
| Pf_16 | 2613 | 36 | 0.60 | 6.00 | **0.18** | 0.00 | 0.56 |
| Pf_17 | 2161 | 36 | 0.60 | 5.93 | **0.16** | 0.00 | 0.50 |
| Pf_18 | 1641 | 39 | 0.63 | 6.82 | **0.22** | 0.07 | 0.63 |
| Pf_19 | 2776 | 34 | 0.60 | 6.37 | **0.17** | 0.06 | 0.46 |
| Pf_20 | 2117 | 37 | 0.65 | 6.73 | **0.13** | 0.00 | 0.46 |
| Pf_21 | 1753 | 35 | 0.61 | 6.19 | **0.08** | 0.00 | 0.31 |
| Pf_22 | 1815 | 34 | 0.62 | 6.53 | **0.21** | 0.02 | 0.75 |
| Pf_23 | 2056 | 36 | 0.61 | 6.69 | **0.18** | 0.01 | 0.61 |
| Pf_24 | 1779 | 36 | 0.64 | 6.81 | **0.11** | 0.03 | 0.33 |

### Estimation of effective and census population sizes

N_e_ ranged from 25 to 228 individuals, the mean N_e_ being 86 individuals and the median N_e_ being 58 individuals (Fig. 2).

**Fig. 2.**
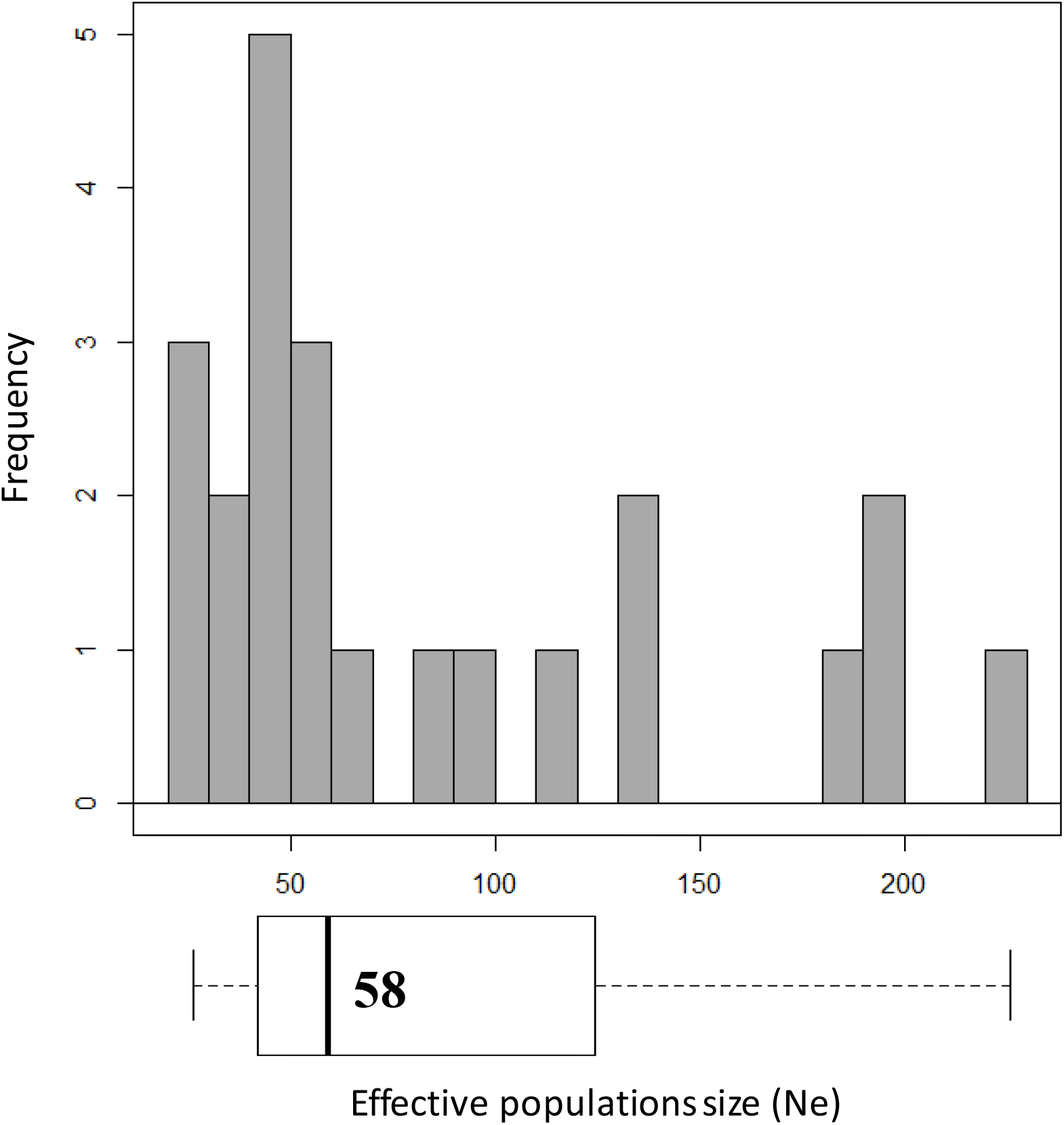
Histogram showing the distribution of the independent effective population sizes estimated using the Wang’s method. The median N_e_ is indicated directly onto the box plot below the histogram.

There was a marginally significant effect of initial populations, differing by the proportion of cysts coming from the different Peruvian populations (*F*_6,16_ = 2.9; *P* = 0.041), but the comparison of means, performed with the Tukey test, was not able to identify distinct homogenous groups. Moreover, there was no significant correlation between mean N_e_ (calculated for each pair of initial and final populations) and the number of cysts coming from each of the four Peruvian populations (data not shown).

The number of newly formed cysts ranged from 1,060 to 3,407 with a mean (±sem) of 2,189 (±116). There was no correlation between N_e_ estimates and the number of newly formed cysts (Pearson’s coefficient cor = -0.18; *P* = 0.41). The number of larvae per cyst, scored for seven final populations, ranged from 196 to 288 with a mean (±sem) of 235 (±12), and a oneway ANOVA showed no significant difference for the number of larvae per cyst between those seven final populations (*F*_6,77_ = 0.74; *P* = 0.62; Fig. 3). Consequently, our estimation of the final census size (N) was 514,415 larvae (2,189 newly formed cysts * 235 larvae per cyst).

**Fig. 3.**
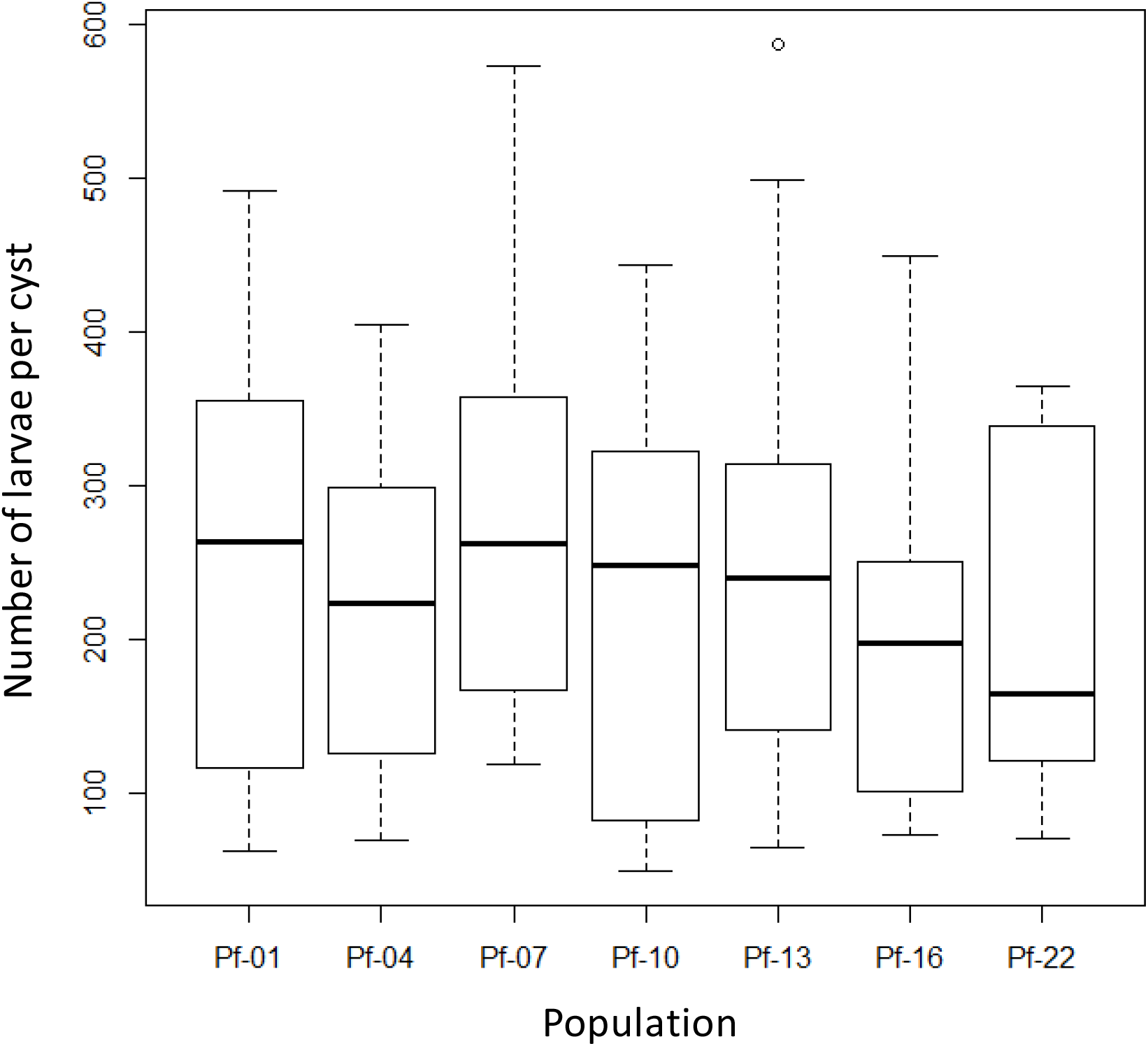
Number of larvae per cyst for the seven final *G. pallida* populations. No significant difference was observed between those populations (F_6,77_ = 0.74; *P* = 0.62).

### Exploring the link between N_e_ and resistance durability

Fixing the initial frequency of an allele at 1%, results showed that for N_e_ = 58, the fixation probability was higher than 50% only for selection coefficients above 0.3, whereas for N_e_ two, five-, 10- or 50-times higher, this threshold of 50% was crossed for lower selection coefficients (Fig. 4).

**Fig. 4.**
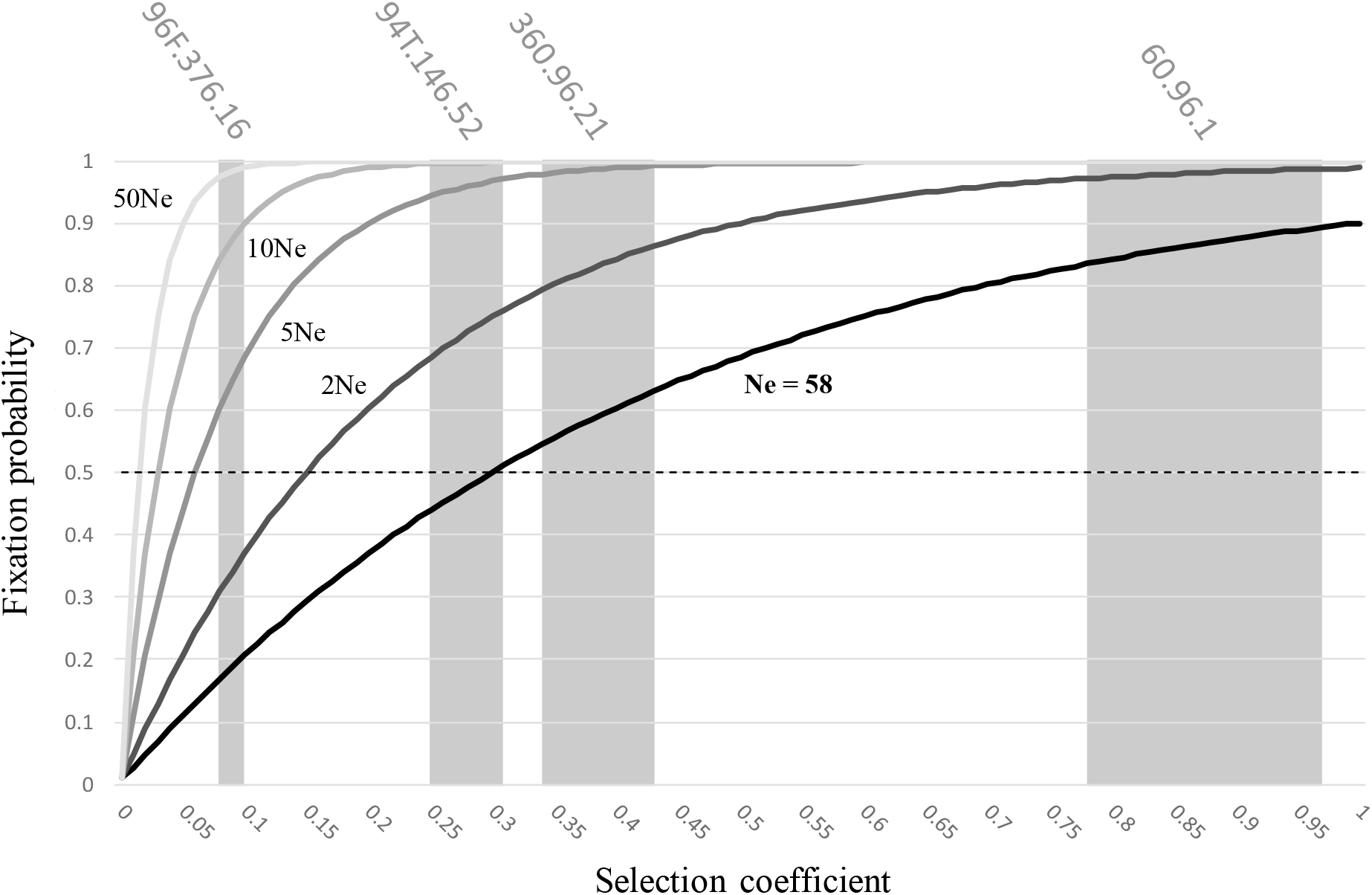
Probability of fixation of an allele (P(A)), with an initial frequency of 1%, according to its selection coefficient (s) for a diploid population showing an effective population size (N_e_) of 58 individuals and for populations showing 2N_e_ (116 individuals), 5N_e_ (290 individuals), 10N_e_ (580 individuals) and 50N_e_ (1,160 individuals). The dotted line indicates the probability of fixation of 50%. Credible intervals for selection coefficients are given as grey backgrounds for four different potato cultivars studied by Fournet et al. (2013). See text for more details.

Using the breeder’s equation, the estimation of the selection coefficients *S* exerted by the four resistant potato genotypes used by Fournet *et al*. (2013) showed that, for N_e_ = 58, the fixation probability was small for the genotype 96F.376.16 (0.08 < *S* < 0.10), medium for 94T.146.52 (0.25 < *S* < 0.31) and 360.96.21 (0.34 < *S* < 0.43) and very high for 60.96.1 (0.78 < *S* < 0.97) (Fig. 4).

## Discussion

This report evaluates the effective size of populations of *Globodera pallida*, the potato cyst nematode. Rather than working with natural populations, which could sometimes harbor very low genetic diversity leading to infinite estimated Ne, we decided here to work with artificial *G. pallida* populations. As expected, and because we have mixed four Peruvian populations that are genetically differentiated, heterozygote deficits observed for initial populations were very high (mean *F*_IS_ = 0.39) and due to both consanguinity and sub-structure, whereas after one generation of mating, heterozygote deficits observed for final populations were lower (mean *F*_IS_ = 0.13) and only due to consanguinity, as in natural *G. pallida* populations (Montarry *et al*., 2015). Moreover, temporal methods assume neither migration nor selection and that the variation in allele frequencies between the samples is only due to genetic drift. Working with artificial population is thus a way to ensure the absence of migration, and allows to reduce the action of selection by using a susceptible potato cultivar. The requirement of an estimation of the generation number leads to difficulties in the evaluation of N_e_ for several species. For example, estimation of N_e_ during infection cycle of plant virus populations is quite complicated because of the lack of estimates of generation times for viruses (Fabre *et al*., 2014). Regarding the beet cyst nematode *H. schachtii*, which is a plurivoltine species, Jan *et al*. (2016) have used two extreme estimations of the generation number. We have not that problem using the monovoltine species *G. pallida* because it performed only one generation over the experiment.

Using the likelihood-based method developed by Wang (2001), the median of the 24 N_e_ estimates was 58 individuals. The census size N of the initial populations was 13,200 individuals (100 cysts * 132 larvae per cyst) and our estimation of the census size of the final populations was 514,415 larvae (2,189 newly formed cysts * 235 larvae per cyst). To obtain the N_e_/N ratio, we computed the harmonic mean of these two values for N as recommended by Waples (2005), yielding 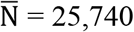, and thus Ne/N ≈ 2.10^-3^. Based on a meta-analysis, values of N_e_/N average only 10–15% (Frankham, 1995; Palstra & Ruzzante, 2008). Thus, effective population sizes are substantially lower than census sizes. For example, the threatened winter run of chinook salmon in the Sacramento River of California has about 2,000 adults, but its effective size was estimated to be only 85 (N_e_/N = 0.04 - Bartley et al. 1992). N_e_ is thus commonly lower than N but in our case the N_e_/N ratio is extremely low, close to values recorded in marine fishes (Hoarau *et al*., 2005).

The effective size of a population is the size of an ideal population which would drift at the same rate as the observed population (Wright, 1931) and all characteristics that deviate between an ideal population and the real populations will cause the effective size (N_e_) to differ from the number of individuals in the population (N). As mentioned above, some of those characteristics are similar between an ideal population and our nematode populations (i.e. no migration, no selection, no possibility for variation of N over generations and no overlapping generations) but others differ. Particularly, our real populations deviate in structure from the assumptions of the ideal population by having unequal sex-ratios and showing non random union of gametes. When larvae of different *G. pallida* populations were inoculated to susceptible potato roots in Petri dishes, the percentage of female produced was on average 60% (Fournet *et al*., 2013). A meta-analysis showed that unequal sex-ratios have modest effects in reducing effective population sizes below actual sizes, resulting in an average reduction of 36% (Frankham, 1995). *G. pallida* populations are characterized by high levels of inbreeding, highlighted here for artificial populations (i.e. *F*_IS_ significantly higher than zero due to consanguinity) and previously highlighted for natural populations (Montarry *et al*., 2015), which could also reduce effective population sizes (Charlesworth, 2009). While random mating generally sustains effective population sizes of pathogens (Barrett *et al*., 2008), inbreeding increases the extent of genetic drift in pathogen populations, resulting in reduced N_e_ (Nunney & Luck, 1988). This factor on its own is however not able to explain the extremely low Ne/N ratio we observe as inbreeding can reduce effective population size by 50% at most (Charlesworth, 2009). The census size of our initial and final nematode populations has increased from 13,200 to 514,415 individuals (i.e. multiplied by 39), indicating clearly that there were several offspring produced per adult. Whether all adults of the initial populations contributed equally to the final populations is however unlikely. It has been documented in cyst nematode species of the genus *Heterodera* that both males and females mate several times, with males contributing differently to the pool of larvae (Green *et al*., 1970; Triantaphyllou & Esbenshade, 1990). Patterns of mitochondrial gene diversity between larvae from the same cyst in a species in which mitochondrial DNA is biparentally transmitted (Hoolahan *et al*., 2011) support the same mating pattern for *G. pallida* (J. Ferreira de Carvalho, S. Fournet & E. J. Petit, unpublished). Using our experimental data, we estimated the variance in family sizes to range between 130 and 2,100 (see Supplementary Table S1), suggesting indeed that some individuals do not contribute at all to the next generation (Hedrick, 2005). Because we estimated N_e_ and N from one generation of J2 larvae to the next, these extreme figures combine both a low probability for each larvae to reach the adult stage, and a high variance in reproductive success for adults. The probability to reach adulthood can here be estimated from the ratio of twice the number of formed cysts (assuming a balanced sex-ratio) to the number of inoculated larvae, that is 2*2,189 / 13,200 = 0.33, meaning than at least 1/3 of all individuals have zero breeding success. Taking into account this proportion of non-breeders is however far from being able to explain the low N_e_/N on its own (see Eq. 5c in Hedrick, 2005). Ultimately, it is the combined impact of all these factors (i.e. extreme variance in family sizes, unequal sex-ratios, and inbreeding) which could explain that the N_e_/N ratio is extremely low in *Globodera pallida* populations.

The low effective population size highlighted here for the potato cyst nematode *G. pallida* is consistent with estimations performed for wild populations of the beet cyst nematode *H. schachtii*: N_e_ around 85 individuals with a N_e_/N ratio less than 1% (Jan *et al*., 2016). Note that because the genetic of *G. pallida* populations deviates from that of an ideal population, the effective population size well describes the genetic drift intensity but estimates poorly the number of individuals that pass on their genes through generations. That explains why we obtained more females (2,189 newly formed cysts) than the estimated N_e_ (58 individuals). It however appears that the effective population size of phytoparasitic cyst nematodes is lower than N_e_ estimates of the free living nematode *Caenorhabditis elegans* (Barrière & Felix, 2005; Sivasundar & Hey, 2005; Cutter, 2006) and of animal parasitic nematodes (e.g. for *Trichostrongylus axei* – Archie & Ezenwa, 2011).

The consequences of a low N_e_ could be important for the control of phytoparasitic cyst nematodes. When N_e_ is large, competition between individuals is fully acting, with no or little interference of random processes, and selection shapes the genetic composition of populations. Conversely, in populations with a small Ne, genetic drift, resulting in stochastic sampling of individuals that will engender the next generation, is prevalent and counters the effect of selection. Consequently, *G. pallida* populations will have a low capacity to adapt to changing environments unless selection intensity is very strong and that could be greatly beneficial for long-term use of plant resistances (McDonald & Linde, 2002). In this paper, for the first time, we were able to determine a selection pressure threshold that allows a better risk assessment and to demonstrate that durable resistance to cyst nematodes is a truly achievable goal. Indeed, for N_e_ = 58, our results showed that the fixation probability was small for the resistant potato genotype 96F.376.16, medium for 94T.146.52 and 360.96.21 and very high for 60.96.1, showing that resistance durability could be anticipated, and that the resistance of cultivars exerting a low selection pressure on pathogen population would be durable only for pathogens showing a low Ne. It is however important to consider that in natural field populations of *G. pallida*, genotype flow have been reported (Picard *et al*., 2004), and those genotype flow could partly compensate the impact of genetic drift (Palstra & Ruzzante, 2008). In cyst nematode, genotype flow has mainly been attributed to the passive transport of cyst through agricultural practices (Alenda *et al*., 2014). Therefore, all agricultural management strategies that reduce genotype flow and thus promote small effective population sizes would be beneficial for the durability of plant resistances.

## Acknowledgments

We gratefully acknowledge Christophe Piriou for his technical help to count the number of cysts and the number of larvae per cyst. Drs. ML Pilet-Nayel and C Lavaud are acknowledged for useful discussions about the breeder’s equation.

## Author contributions

SBV, RM, and JM performed the experiments according to a protocol elaborated jointly by SF, EG and JM. PLJ, EJP and JM analyzed the data. SF, EJP, EG and JM wrote the text and prepared the figures. All authors edited the paper and have approved the current version.

## Data archival location

All data used in this article are available at datadryad.org (doi: to be completed).

## Supporting Information

Additional supporting information may be found on the online version of this article.

**Fig. S1.**
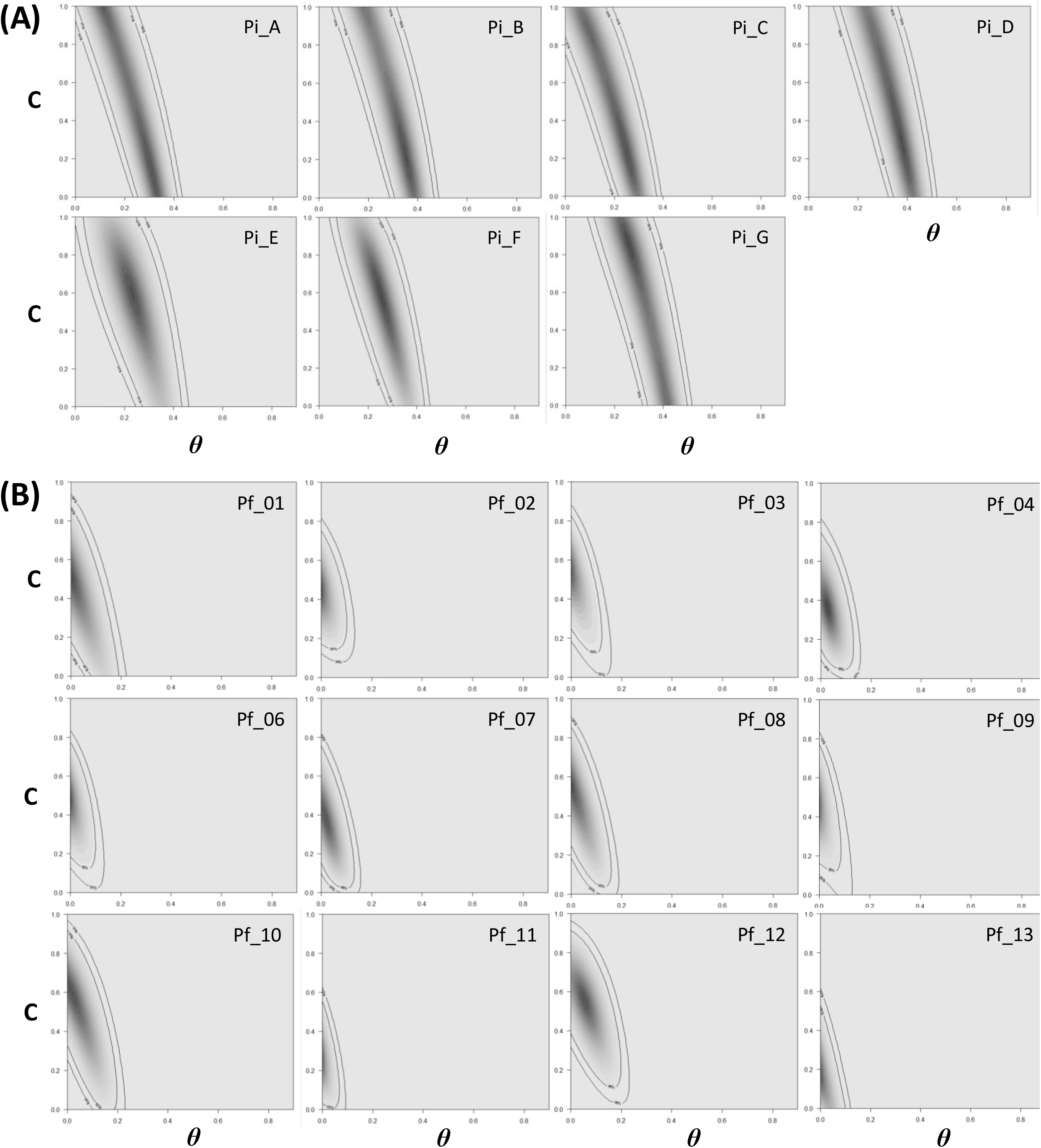
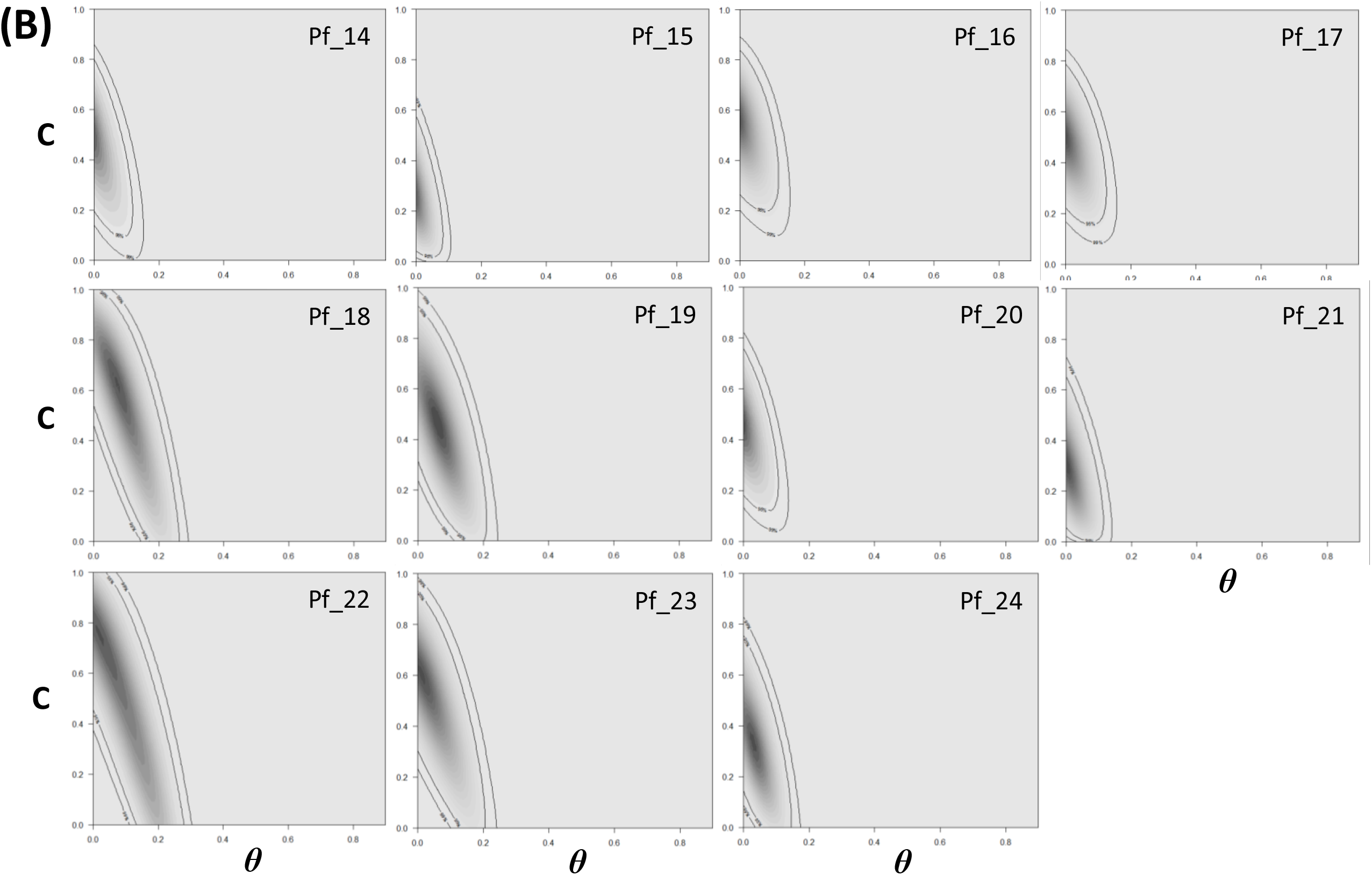
Likelihood surfaces showing estimated *ϑ* and C values with 95 and 99% confidence envelopes (internal and external envelopes of the highest likelihoods, visualized as grey shades, respectively) for (**A**) the seven initial *G. pallida* populations, (**B**) the 23 final *G. pallida* populations showing significant heterozygote deficits. *ϑ* and C are represented on the x-axis and y-axis, respectively.

**Table S1.** Estimation of the variance in family size (S^2^_k_) from the effective population sizes (N_e_) and proportions of inbred matings (C) estimated for each final population. The computation was not possible for Pf_15 (no N_e_ estimate). The estimation was based on Caballero and Hill (1992, Eq. 10), with α computed from C after Ghai (1969, Eq. 17), and N = 13,200, the census size of initial populations (see Text).

| Final populations | C | $\alpha$ | $N_e$ | $S^2_k$ |
| --- | --- | --- | --- | --- |
| Pf_01 | 0.52 | 0.213 | 44.59 | **721** |
| Pf_02 | 0.45 | 0.170 | 59.90 | **583** |
| Pf_03 | 0.54 | 0.227 | 41.79 | **751** |
| Pf_04 | 0.35 | 0.119 | 30.95 | **1257** |
| Pf_05 | 0.00 | 0.000 | 24.86 | **2122** |
| Pf_06 | 0.48 | 0.188 | 40.34 | **837** |
| Pf_07 | 0.38 | 0.133 | 112.09 | **336** |
| Pf_08 | 0.56 | 0.241 | 136.54 | **223** |
| Pf_09 | 0.46 | 0.176 | 94.34 | **366** |
| Pf_10 | 0.59 | 0.265 | 193.49 | **151** |
| Pf_11 | 0.25 | 0.077 | 86.50 | **494** |
| Pf_12 | 0.57 | 0.249 | 227.76 | **132** |
| Pf_13 | 0.18 | 0.052 | 136.63 | **333** |
| Pf_14 | 0.49 | 0.194 | 34.48 | **968** |
| Pf_15 | 0.26 | 0.081 | / |  |
| Pf_16 | 0.56 | 0.241 | 27.06 | **1131** |
| Pf_17 | 0.50 | 0.200 | 28.36 | **1163** |
| Pf_18 | 0.63 | 0.299 | 47.05 | **591** |
| Pf_19 | 0.46 | 0.176 | 58.41 | **591** |
| Pf_20 | 0.46 | 0.176 | 187.01 | **184** |
| Pf_21 | 0.31 | 0.101 | 48.26 | **838** |
| Pf_22 | 0.75 | 0.429 | 56.58 | **408** |
| Pf_23 | 0.61 | 0.281 | 63.56 | **450** |
| Pf_24 | 0.33 | 0.110 | 195.37 | **202** |

## References

Adams HH, Osborne WW, Webber AJ. 1982. Effect of temperature on development and reproduction of Globodera solanacearum. Nematropica 12: 305–311.

Alenda, C, Montarry J, Grenier E. 2014. Human influence on the dispersal and genetic structure of French Globodera tabacum populations. Infection. Genetics and Evolution 27: 309–317.

Araki H., Waples RS, Blouin MS. 2007. A potential bias in the temporal method for estimating Ne in admixed populations under natural selection. Molecular Ecology 16: 2261–2271.

Archie EA, Ezenwa VO. 2011. Population genetic structure and history of a generalist parasite infecting multiple sympatric host species. International Journal of Parasitology 41: 89–98.

Barrett LG, Thrall PH, Burdon JJ, Linde CC. 2008. Life history determines genetic structure and evolutionary potential of host–parasite interactions. Trends in Ecology and Evolution 23: 678–685.

BarrièreA. Félix MA. 2005. High local genetic diversity and low outcrossing rate in Caenorhabditis elegans natural populations. Current Biology 15: 1176–1184.

Bartley D, Bagley M, Gall G, Bentley B. 1992. Use of linkage disequilibrium data to estimate effective size of hatchery and natural fish populations. Conservation Biology 6: 365–375.

Belkhir K, Borsa P, Chikhi L, Raufaste N, Bonhomme F. 1996–2004. GENETIX 4.05, logiciel sous Windows TM pour la génétique des populations. Laboratoire Génome, Populations, Interactions, CNRS UMR 5000, Université de Montpellier II, Montpellier (France).

Betancourt M, Fereres A, Fraile A, García-Arenal F. 2008. Estimation of the effective number of founders that initiate an infection after aphid transmission of a multipartite plant virus. Journal of Virology 82: 12416–12421.

Boucher AC, Mimee B, Montarry J, Bardou-Valette S, Bélair G, Moffett P, Grenier E. 2013. Genetic diversity of the golden potato cyst nematode Globodera rostochiensis and determination of the origin of populations in Quebec, Canada. Molecular Phylogenetics and Evolution 69: 75–82.

Charlesworth B. 2009. Effective population size and patterns of molecular evolution and variation. Nature Reviews Genetics 10: 195–205.

Cotton JA, Lilley CJ, Jones LM, Kikuchi T, Reid AJ, Thorpe P, Tsai IJ, Beasley H, Blok V, Cock PJA et al. 2014. The genome and life-stage specific transcriptomes of Globodera pallida elucidate key aspects of plant parasitism by a cyst nematode. Genome Biology 15: R43.

Criscione CD, Blouin MS. 2005. Effective sizes of macroparasite populations: a conceptual model. Trends in Parasitology 21: 212–217.

Crow JF, Kimura M. 1970. An introduction to population genetics theory. New York, USA: Harper and Row.

Crow JF, Kimura M. 1972. The effective number of a population with overlapping generations: a correction and further discussion. The American Journal of Human Genetics 24: 1–10.

Cutter AD. 2006. Nucleotide polymorphism and linkage disequilibrium in wild populations of the partial selfer Caenorhabditis elegans. Genetics 172: 171–184.

Damgaard C, Giese H. 1996. Genetic variation in Danish populations of Erysiphe graminis f.sp. hordei: Estimation of gene diversity and effective population size using RFLP data. Plant Pathology 45: 691–696.

Duan X, Tellier A, Wan A, Leconte M, de Vallavielle-Pope C, Enjalbert J. 2010. Puccinia striiformis f.sp tritici presents high diversity and recombination in the over-summering zone of Gansu, China. Mycologia 102: 44–53.

Fabre F, Montarry J, Coville J, Senoussi R, Simon V, Moury B. 2012. Modelling the evolutionary dynamics of viruses within their hosts: a case study using high-throughput sequencing. PLoS Pathogens 8: e1002654.

Fabre F, Moury B, Johansen EI, Simon V, Jacquemond M, Senoussi R. 2014. Narrow bottlenecks affect Pea seedborne mosaic virus populations during vertical transmission but not during leaf colonization. PLoS Pathogens. 10: e1003833.

Fournet S, Kerlan MC, Renault L, Dantec JP, Rouaux C, Montarry J. 2013. Selection of nematodes by resistant plants has implications for local adaptation and cross-virulence: local adaptation and cross-virulence in Globodera pallida. Plant Pathology 62: 184–193.

Frankham R. 1995. Effective population size / adult population size ratios in wildlife: a review. Genetics Research 66: 95–107.

Frankham R, Ballou JD, Briscoe DA. 2002. Introduction to conservation genetics. Cambridge, UK: Cambridge University Press.

Fraser AS. 1972. An introduction to population genetic theory. In: Crow JF, Kimura M, eds. Teratology, Vol. 5. New York, USA: Harper and Row, 386–387.

Gandon S, Michalakis Y. 2002. Local adaptation, evolutionary potential and host–parasite coevolution: interactions between migration, mutation, population size and generation time. Journal of Evolutionary Biology 15: 451–462.

Gherman A, Chen PE, Teslovich TM, Stankiewicz P, Withers M, Kashuk CS, Chakravarti A, Lupski JR, Cutler DJ, Katsanis N. 2007. Population bottlenecks as a potential major shaping force of human genome architecture. PLoS Genetics 3: 1223–1231.

Green CD, Greet DN, Jones FGW. 1970. The influence of multiple mating on the reproduction and genetics of Heterodera rostochiensis and H. schachtii. Nematologica 16: 309–326.

Grenier E, Fournet S, Petit E, Anthoine G. 2010. A cyst nematode ‘species factory’ called the Andes. Nematology 12: 163–169.

Gutiérrez S, Michalakis Y, Blanc S. 2012. Virus population bottlenecks during within-host progression and host-to-host transmission. Current Opinion in Virology 2: 1–10.

Hedrick P. 2005. Large variance in reproductive success and the Ne/N ratio. Evolution 59: 1596–1599.

Hijmans RJ, Spooner DM. 2001. Geographic distribution of wild potato species. American Journal of Botany 88: 2101–2112.

Hoarau G, Boon E, Jongma DN, Ferber S, Palsson J, der Veer HWV, Rijnsdorp AD, Stam WT, Olsen JL. 2005. Low effective population size and evidence for inbreeding in an overexploited flatfish, plaice (Pleuronectes platessa L.). Proceedings of the Royal Society of London B: Biological Sciences 272: 497–503.

Holleley CE, Nichols RA, Whitehead MR, Adamack AT, Gunn MR, Sherwin WB. 2014. Testing single-sample estimators of effective population size in genetically structured populations. Conservation Genetics 15: 23–35.

Hoolahan AH, Blok VC, Gibson T, Dowton M. 2011. Paternal leakage of mitochondrial DNA in experimental crosses of populations of the potato cyst nematode Globodera pallida. Genetica 139: 1509–1519.

Jan PL, Gracianne C, Fournet S, Olivier E, Arnaud JF, Porte C, Bardou-Valette S, Denis MC, Petit EJ. 2016. Temporal sampling helps unravel the genetic structure of naturally occurring populations of a phytoparasitic nematode. 1 Insights from the estimation of effective population sizes. Evolutionary Applications 9: 489–501.

Johnson JA, Bellinger MR, Toepfer JE, Dunn P. 2004. Temporal changes in allele frequencies and low effective population size in greater prairie-chickens. Molecular Ecology 13: 2617–2630.

Johnson R. 1981. Durable resistance: definition of genetic control and attainment in plant breeding. Phytopathology 71: 567–568.

Johnson R. 1984. A critical analysis of durable resistance. Annual Review of Phytopathology 22: 309–330.

Jones MGK, Northcote DH. 1972. Nematode-induced syncytium - A multinucleate transfer cell. Journal of Cell Science 10: 789–809.

Kimura M. 1962. On the probability of fixation of mutant genes in a population. Genetics 47: 713–719.

Kimura M, Maruyama T, Crow JF. 1963. The mutation load in small populations. Genetics 48:1303–1312.

Langella O. 2000. POPULATIONS 1·2: population genetic software, individuals or population distance, phylogenetic trees. http://bioinformatics.org/~tryphon/populations.

Luikart G, Ryman N, Tallmon DA, Schwartz MK, Allendorf FW. 2010. Estimation of census and effective population sizes: the increasing usefulness of DNA-based approaches. Conservation Genetics 11: 355–373.

Lush JL. 1937. Animal breeding plans. Ames, IA, USA: Iowa State College Press.

McDonald BA, Linde C. 2002. Pathogen population genetics, evolutionary potential, and durable resistance. Annual Review of Phytopathology 40: 349–379.

Monsion B, Froissart R, Michalakis Y, Blanc S. 2008. Large bottleneck size in Cauliflower mosaic virus populations during host plant colonization. PLoS Pathogens 4: e1000174.

Montarry J, Jan PL, Gracianne C, Overall ADJ, Bardou-Valette S, Olivier E, Fournet S, Grenier E, Petit EJ. 2015. Heterozygote deficits in cyst plant-parasitic nematodes: possible causes and consequences. Molecular Ecology 24: 1654–1667.

Nei M. 1978. Estimation of average heterozygosity and genetic distance from a small number of individuals. Genetics 89: 583–590.

Nei M, Tajima F. 1981. Genetic drift and estimation of effective population size. Genetics 98: 625–640.

Nei M, Maruyama T, Chakaraborty R. 1975. The bottleneck effect and genetic variability in populations. Evolution 29: 1–10.

Nunney L, Luck RF. 1988. Factors influencing the optimum sex ratio in a structured population. Theoretical Population Biology 33: 1–30.

Overall ADJ, Nichols RA. 2001. A method for distinguishing consanguinity and population substructure using multilocus genotype data. Molecular Biology and Evolution 18: 2048–2056.

Palstra FP, Ruzzante DE. 2008. Genetic estimates of contemporary effective population size: what can they tell us about the importance of genetic stochasticity for wild population persistence? Molecular Ecology 17: 3428–3447.

Picard D, Plantard O, Scurrah M, Mugniéry D. 2004. Inbreeding and population structure of the potato cyst nematode (Globodera pallida) in its native area (Peru). Molecular Ecology 13: 2899–2908.

Plantard O, Porte C. 2004. Population genetic structure of the sugar beet cyst nematode Heterodera schachtii: a gonochoristic and amphimictic species with highly inbred but weakly differentiated populations. Molecular Ecology 13: 33–41.

Pudovkin AI, Zaykin DV, Hedgecock D. 1996. On the potential for estimating the effective number of breeders from heterozygote-excess in progeny. Genetics 144: 383–387.

R Core Team. 2014. R: a language and environment for statistical computing. R Foundation for statistical computing, Vienna, Austria. https://www.R-project.org.

Sasser JN, Freckman DW. 1987. A world perspective on nematology: the role of the society. In: Veech JA, Dickson DW, eds. Vistas on nematology. Hyatssville, Maryland, USA: Society of Nematologists, 7–14.

Sivasundar A, Hey J. 2005. Sampling from natural populations with RNAi reveals high outcrossing and population structure in Caenorhabditis elegans. Current Biology 15: 1598–1602.

Sobczak M, Golinowski W. 2011. Cyst nematodes and syncytia. In: Jones JT, GheysenG, Fenoil C, eds. Genomics and molecular genetics of plant-nematode interactions. Dordrecht, The Netherlands: Springer, 61–82.

Stukenbrock EH, Bataillon T, Dutheil JY, Hansen TT, Li R, Zala M, McDonald BA, Wang J, Schierup MH. 2011. The making of a new pathogen: insights from comparative population genomics of the domesticated wheat pathogen Mycosphaerella graminicola and its wild sister species. Genome Research 21: 2157–2166.

Tajima F, Nei M. 1984. Note on genetic drift and estimation of effective population size. Genetics 106: 569–574.

Tallmon DA, Koyuk A, Luikart G, Beaumont MA. 2008. Onesamp: a program to estimate effective population size using approximate bayesian computation. Molecular Ecology Resources 8: 299–301.

Tellier A, Brown JKM. 2007. Stability of genetic polymorphism in host-parasite interactions. Proceedings of the Royal Society of London B: Biological Sciences 274: 809–817.

Tellier A, Brown JKM. 2009. The influence of perenniality and seed banks on polymorphism in plant-parasite interactions. American Naturalist 174: 769–779.

Triantaphyllou AC, Esbenshade PR. 1990. Demonstration of multiple mating in Heterodera glycines with biochemical markers. Journal of Nematology 22: 452–456.

Vanderplank JE. 1963. Disease resistance in plants. New York, USA: Academic Press.

Wang J. 2001. A pseudo-likelihood method for estimating effective population size from temporally spaced samples. Genetics Research 78: 243–257.

Wang J. 2005. Estimation of effective population sizes from data on genetic markers. Philosophical Transactions of the Royal Society of London B – Biological Sciences 360: 1395–1409.

Wang J, Whitlock MC. 2003. Estimating effective population size and migration rates from genetic samples over space and time. Genetics 163: 429–446.

Waples RS. 2005. Genetic estimates of contemporary effective population size: to what time periods do the estimates apply? Molecular Ecology 14: 3335–3352.

Waples RS, Do CHI. 2008. LDNE: a program for estimating effective population size from data on linkage disequilibrium. Molecular Ecology Resources 8: 753–756.

Waples RS, Do CHI. 2010. Linkage disequilibrium estimates of contemporary Ne using highly variable genetic markers: an untapped resource for applied conservation and evolution. Evolutionary Applications 3: 244–262.

Wright S. 1931. Evolution in Mendelian populations. Genetics 16: 97–159.

Zhan J, Mundt CC, McDonald BA. 2001. Using restriction fragment length polymorphisms to assess temporal variation and estimate the number of ascospores that initiate epidemics in field populations of Mycosphaerella graminicola. Phytopathology 91: 1011–1017.

Zhdanova OL, Pudovkin AI. 2008. Nb_HetEx: a program to estimate the effective number of breeders. Journal of Heredity 99: 694–695.

Zwart MP, Daròs JA, Elena SF. 2011. One is enough: In vivo effective population size is dose-dependent for a plant RNA virus. PLoS Pathogens 7: e1002122.

## References

Caballero A, Hill WG. 1992. Effective size of nonrandom mating populations. Genetics 130: 909–916.

Ghai GL. 1969. Structure of populations under mixed random and sib mating. Theoretical and Applied Genetics 39: 179–182.

